# Graphing and reporting heterogeneous treatment effects through reference classes

**DOI:** 10.1101/708958

**Authors:** James A. Watson, Chris C. Holmes

## Abstract

**Background:** Exploration and modelling of individual treatment effects and treatment heterogeneity is an important aspect of precision medicine in randomized controlled trials (RCTs). The usual approach is to develop a predictive model for individual outcomes and then look for an interaction effect between treatment allocation and important patient covariates. However, such models are prone to overfitting and multiple testing, and typically demand a transformation of the outcome measurement, for example, from the absolute risk in the original RCT to log-odds of risk in the predictive model.

**Methods:** We show how reference classes derived from background information can be used to alleviate this problem through a two-stage approach where we first estimate a key aspect of heterogeneity in the trial population and then explore for an interaction with the treatment effect along this axis of variation. This bypasses the search for interactions, protecting against multiple testing, and allows for exploration of heterogeneous treatment effects on the original outcome scale of the RCT. This would typically be applied to multivariate models of baseline risk to assess the stability of average treatment effects with respect to the distribution of risk in the population studied. We show how ‘local’ and ‘tilting’ schemes based on ranking patients by baseline risk can be used as a general approach for exploring heterogeneity of treatment effect.

**Results:** We illustrate this approach using the single largest randomised treatment trial in severe falciparum malaria and show how the estimated treatment effect in terms of absolute mortality risk reduction increases considerably for higher risk strata.

## Background

Randomised controlled trials (RCTs) provide the best causal evidence base for estimating the population average treatment effect (ATE) for a given intervention. This can then be used to support optimal decision making at the population level. At the individual level, however, the ATE is often unrepresentative of the true individual treatment effect (ITE) for a large proportion of patients. This may arise when the absolute treatment effect varies as a function of the individual risk of a negative outcome (e.g. treatment failure, death, severe adverse event, etc.). It has been previously shown that the baseline risk of a negative outcome is often highly skewed in patient populations for many diseases and conditions [1]. Hence the average risk, on which the ATE is estimated, may not be a good summary of the individual risk [2].

Due to this heterogeneity in baseline risk, guidelines for the reporting and assessment of RCT results have recommended the use of multivariate risk prediction tools for patient stratification [3, 4]. Baseline predicted risk provides a single dimension over which individuals can be compared and average outcomes assessed. This provides a methodology for risk-based reference class forecasting and a principled way of assessing personalised and heterogeneous treatment effects (HTE) [5].

The assessment of HTEs, and thus the stability of trial results, has classically been done by constructing parametric models targeting the ITE, conditioning on either the baseline risk or key variables predictive of the baseline risk [6]. However, estimating an ITE brings with it two challenges, one foundational and one practical. The foundational challenge is that the estimand of an ITE is counterfactual, targeting the expected difference between an observed (actual) outcome and an unobserved (potential) outcome that would have occurred should the individual have been given an alternative treatment. The practical challenge is that statistical methods targeting ITEs invariably need to transform the outcome measurement to allow for contextual modelling, such as transforming to the log-odds scale under logistic regression, or proportional hazards for time-to-event data. These transformations inevitably make results dependent on parametric model assumptions and link functions concerning the effect of patient’s baseline covariates on their outcomes. Moreover, when testing for evidence of clinically significant variation in the ITE, considerable care must be taken not to overfit to the data especially when considering a large number of potential predictor variables [7]

Here we promote a non-parametric (model-free) approach to the estimation and assessment of HTEs using risk-based reference class forecasting as our example. A reference class, constructed through a sample re-weighting scheme, can be used to explore for treatment effect variation in a target population different to the one collected through the RCT. A targeted treatment effect can then be estimated for this new population. In particular we consider reference classes of individuals such that the distributions of baseline risk are different to the population sampled under the RCT.

These reference classes can then be used for two purposes. Firstly, to estimate counterfactual ITEs. This can be done through ‘local’ reference classes centred around an individual of interest. Secondly, to estimate an average treatment effects in ‘tilted’ populations with different risk distributions to that of the RCT. We denote these as tilted average treatment effects (TATEs). The latter may be particularly useful if the risk profile of the trial population systematically under, or over, estimates the profile of risk of the population for which the intervention is aimed at. The TATE can then be used to explore stability of the RCT outcome with respect to this variation in population risk. We show how these different re-weighting schemes correspond to different bias-variance trade-offs for the reference class estimator, and we provide guidelines on graphing of results. It is important to note that estimands from reference classes target the same units of treatment effect as that defined in the original RCT. This allows for direct comparison between estimates. This is not true in general for HTE models, which, as noted above, may have to transform the outcome prior to modelling, for example transforming an absolute effect to the log-odds scale for a binary response. We note that providing individualised predictions through local-weighted averaging has a rich history in the field of kernel smoothing methods, for the example the Nadaraya-Watson estimator [8, 9, 10].

For illustration we consider the single largest trial of life-saving interventions in severe malaria, demonstrating that the superiority of parenteral artesunate stems from its very large effect in the most severely ill patients.

## Methods

### Multivariate risk based ranking of trial individuals

In the following we consider RCT data of the form 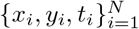 where *x_i_* is a vector of baseline patient covariates for the *i*’th patient, and *y_i_* is their observed outcome after receiving a randomised treatment allocation indicated by *t_i_*. We are interested in pairwise comparisons between *K* treatment arms *t_i_* ∈ {1,…, *K*}.

We assume that it is possible to construct a prior ‘risk mapping’ *Q* : *X* → [0, 1] for the outcome which maps each patient in the trial to their corresponding empirical risk quantile, agnostic of treatment. So that, *Q*(*x_i_*) = 0 denotes the subject least at risk of the negative outcome, and *Q*(*x_j_*) = 1 the subject most at risk. In practice this mapping could be derived by estimating a function *f* : *X* → *Y* using data from a different source (this could include observational data as the risk is agnostic of the treatment received); computing *f* (*x_i_*) for each patient; and then mapping *f* (*x_i_*) onto the empirical risk distribution for the *N* patients in the trial. For almost all major conditions, there will exist external data on which to build this risk-based ranking [1]. If this is not the case, it is also possible to build an internal risk model by ‘retrodiction’: fitting the function *f* to the trial data at hand. Simulation studies suggest that these internal models introduce little bias into the procedure [11].

In the following, for simplicity we assume that the patient index *i* has subsequently been sorted according to the risk prediction with *Q*(*x*_1_) = 0, and *Q*(*x_N_*) = 1. In general, and in absence of ties, 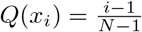.

We can then use the profile mapping to a reference class, in this case risk, constructed prior to the RCT analysis to explore for heterogeneous treatment effects. This removes the need for multiple testing of treatment interactions as it targets the reference class for the exploration of treatment heterogeneity. The use of risk-based reference classes for exploring HTEs have been advocated previously, but limited to quintile or quartile subgroups [3, 4, 12]. We now consider using the risk-ranked individuals to estimate ITEs and TATEs.

### Local smoothing estimation of a risk-based ITE using reference class forecasting

Localised re-weighting kernels generalise the idea of partitioning subjects into quintile or quartile subgroups [8]. Local kernels target a specific individual focused at their quantile of risk *q_i_* for the *i*’th subject by considering the treatment outcome of other individuals in a local neighbourhood of risk-adjacent individuals, with *q*’s close to *q_i_*. These local reference classes are parameterised by their bandwidth (radius) *γ* ∈ [0, 1], which defines the proportion of patients in the window ‘close’ to patient *i*, which are used to estimate the ITE of the *i^th^* patient. This in turn characterises the effective sample size of the reference class forecasting method. The simplest local reference class forecasting weighting scheme uses the ‘boxcar’ function that gives equal weight to all subjects in the local window when estimating the ITE. So for a window of width *γ* we can estimate the ITE as

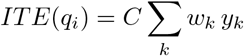

where *C* = (Σ*_k_ w_k_*)^−1^ and *w_k_* = 1 for subjects in the window around subject *i* and *w_k_* = 0 outside of the window, i.e.,

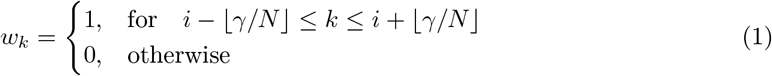

This uses patient data in the risk quantiles of width 2*γ* centred around the *i*’th patient. For *q_i_*’s from the lowest and highest risk patients, for whom there are not ⌊*γ*/*N*⌋ risk-adjacent patients on each side, we take the convention to define the ITE as that of the patients ⌊*γ*/*N*⌋ and *N* − ⌊*γ*/*N*⌋, respectively. This preserves symmetry at the risk ‘tails’. The boxcar kernel is known to be problematic as it varies in a non-smooth way as subjects enter into and leave the kernel, as illustrated in the supplementary Figure S3. A better approach is to use a window that gradually down weights the influence of subjects in the estimate as subjects move away from the prediction point at *q_i_*.

A smoother, improved reference class forecasting method uses the Epanechnikov kernel, again defined on the risk quantiles of radius of width *γ* around the patient *i*, with the weight given as:

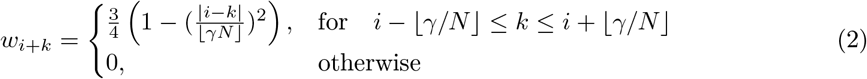

We use the same convention at the edges of the risk distribution, for the patients *i* < ⌊*γ*/*N*⌋ and *i* > *N* − ⌊*γ*/*N*⌋. Under an Epanechnikov reweighting scheme, the weights slowly decay as a function of the distance from the *i*’th datapoint. Both the boxcar and Epanechnikov kernels, centred around the 25% risk-quantile, are illustrated in Figure 1.

**Figure 1:**
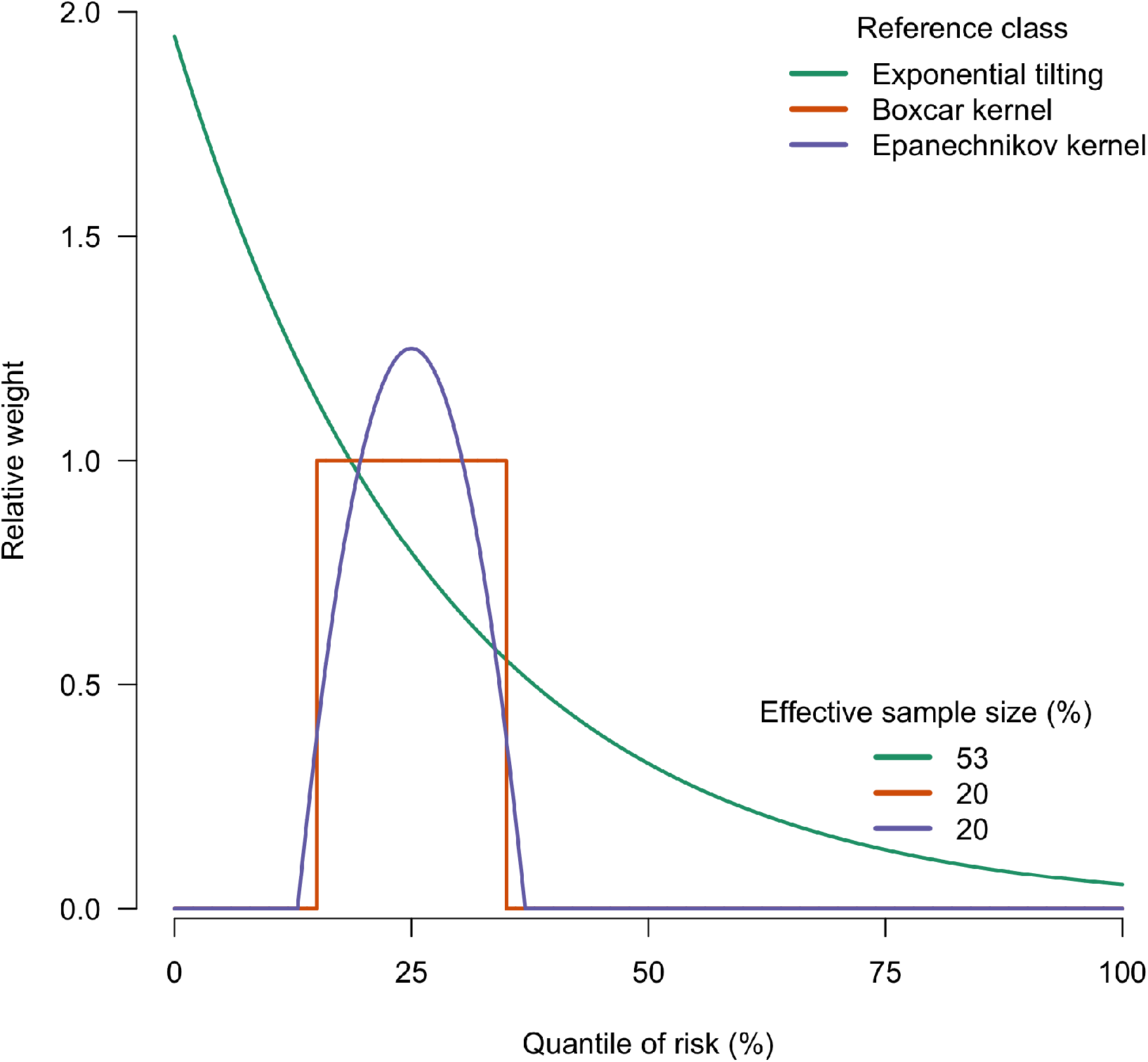
Two local kernel weighting schemes for ITE estimation and one global re-weighting scheme for TATE estimation. The reweighting schemes represented all reduce the effective sample size of the original data and target an average risk corresponding to the 25^*th*^ risk quantile. The scale of the y-axis is chosen so that the sum of the weights is equal to the effective sample size. The effective sample sizes as a percentage of the original data are shown in the legend.

These ‘local’ reference class forecasting methods are symmetrical around the prediction for the patient *i* i.e. they use an equal number of datapoints each side of *i*. However, they both can be adapted so that the bandwidth varies, exploiting the maximum possible information around the patient *i* and preserving symmetry. For example, at the median risk quantile a varying bandwidth method would use all the data. We denote these maximal bandwidth local reference classes, and define the size of the window of information around the *i*’th patient as min(*i, N* − *i*), where the parameter *γ* now specifies the minimal value that this window can take.

### Estimation of a risk-based TATE using reference class forecasting with exponential tilting

Local reweighting schemes provide a principled approach for determining an ITE for a given patient in the trial up to a certain accuracy, with a certain bias-variance trade-off. A different goal is to estimate population average treatment effects but in populations with a different risk distribution to that of the original trial. Often external populations for which the intervention is intended may differ to those of the trial due to issues such as non-representative inclusion criteria, selection bias, or geographical clustering. We denote the estimation of the expected effect under a different population as a conditional average treatment effect (cATE). One interesting, and identifiable, external population can be made through tilting the original sample set through re-weighting the contribution from each sample. Exponential tilting of the population weights is one example. Under this scheme we can consider estimating the average treatment effect in an external tilted population that contains more higher-risk subjects (or more lower risk) as compared with the original trial. In this scheme the *i*’th patient with baseline covariates *x* is attributed a weight proportional to *e*^*λQ*(*x*)^ in the estimate of the external average treatment effect, where the free parameter *λ* determines the overall effective sample of the scheme and how far ‘tilted’ the weights are to the highest risk patients (*λ* > 0) or the lowest risk patients (*λ* < 0). The choice of *λ* = 0 recovers the original ATE. The ratio of the relative weight *w*_1_ (the lowest risk patient) to the relative weight *w_N_* (the highest risk patient) is thus *e^−λ^*. This is akin to estimating the ATE in a population whose participants are recruited with probability *e*^*λQ*(*x*)^ relative to the original trial population.

### Effective sample size and bias-variance trade off

Reference class estimators using re-weighting schemes - whether they are global or local - provide unbiased estimators of the targeted treatment effect in the ‘local’ or ‘tilted’ population, but have increased variance with respect to that of the ATE estimated from the original RCT. This is a consequence of the reduced effective sample size within the reference class. The effective sample size can be thought of as the number of patients (each given weight 1) required to obtain the same accuracy of estimation as in the weighted population. As a function of the weights *w_i_*, the effective sample size is given by: 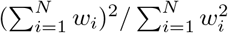. The effective sample size is equal to *N* when all weights *w_i_* are equal to 1 and is strictly less than *N* otherwise. The effective sample size is directly related to the power to detect HTEs using a reweighted reference class. The more distinct the class, the lower the effective sample size, and thus the lower the power to reject the null hypothesis for any given HTE size.

For local reference classes the effective sample size decreases with decreasing bandwidth of the kernel. This relates to a bias-variance trade-off in estimating the ITE at a reference quantile. The more localised the kernel the lower the bias to estimate the target ITE but the greater the standard error of the estimate, which is a function of square-root of the effective sample size. For instance, a kernel that only includes *x_i_* at a reference point has zero-bias for the unique ITE, but infinite variance of the estimate as only one outcome is observed.

### Properties of reweighting schemes under no heterogeneity

It is interesting to note that under an assumption of “no treatment effect heterogeneity”, any weighted average of the outcomes is an unbiased estimator of the ATE albeit with increased variance. If we consider the event “no HTE” as a null-hypothesis then, under this null, the reweighted reference class ITEs will be distributed around the ATE with a variance determined by the effective sample size. This provides for a formal testing framework able to reject this null-hypothesis at a certain level of significance, *α*, should there be HTE under an alternative hypothesis.

## Results

### The HTE of parenteral artesunate for the treament of severe falciparum malaria

Severe falciparum malaria is a medical emergency characterised by potentially lethal vital organ dysfunction. Mortality is high even in presence of effective treatment, and is strongly dependent on the number and the severity of the complications at presentation. Severe malaria does not have a single case definition but represents a spectrum of illness where risk of death is highly predictable from baseline admission covariates. The best prognostic covariates are the presence of coma, concentrations of base deficit and blood urea nitrogen, and total parasite biomass [13]. The operational definition of severe malaria as given by the World Health Organisation (WHO) [14] provides cutoffs that allow for efficient triage of patients at the highest risk of death, and standardisation of clinical studies of novel interventions in this clinically important subgroup. However, due the multifactorial nature of the illness, estimating HTE within this subgroup is well suited for a multivariate risk modelling approach.

A key recent advance in the last decade in the treatment of severe malaria has been the introduction of parenteral artesunate. This has been shown to reduce mortality by up to 30% compared to parenteral quinine [15, 16]. In this section we illustrate our approach to reference class forecasting using the single largest study ever conducted in severe malaria, which compared artesunate to quinine in African children (AQUAMAT) [16]. The results of this trial lead to parenteral artesunate becoming the WHO recommended treatment worldwide for severe falciparum malaria. In endemic countries where there is currently no evidence of clinically significant artemisinin drug resistance (i.e. everywhere except Southeast Asia [17]), there is no *a priori* reason to believe that there are any severe malaria patients who do not benefit from artesunate over quinine. However, characterising heterogeneity in the treatment effect of artesunate is important for understanding the unique pharmacodynamics underlying its superiority. This is especially important in light of emerging resistance to the artemisisin derivatives in Southeast Asia [18]. Characterising the relationship between baseline risk and treatment effect also provides a rational approach for defining study inclusion criteria in order to maximise power at a given sample size.

The following illustrates our suggested approach to assessing HTEs from RCT data and how this heterogeneity should be graphically visualised with the help of a multivariate risk model. We first constructed a multivariate risk model of death from severe malaria using data from over 4000 patients from multiple randomised and observational studies [15, 19, 20, 21, 22, 23, 24, 25]. For all patients in this training dataset, risk of in-hospital mortality was then estimated using a mixed-effects logistic regression model with the presence of coma (yes/no), and the base deficit concentration (mEq/L) as fixed effects. Study code and country of patient recruitment were added as random effect terms. This model was then used to predict the baseline probability of death in all patients (*n* = 5483) recruited in the AQUAMAT study. The assigned risk of outcome was highly predictive of death in the AQUAMAT study (see supplementary Figure S2).

The original publication of the AQUAMAT study results reported no significant effect from a Mantel-Haenszel analysis of the pre-defined subgroups [16]. This is unsurprising as it is well accepted that assessing HTE using single patient covariates lacks power and is prone to false positive results [4, 12]. Using reference class schemes, however, Figure 2 shows that it is visually apparent that treatment effect is strongly dependent on the baseline risk of death. Panel A in Figure 2 shows a standard quintile subgroup plot such as that recommended in [3]. Each subgroup has an effective sample size equal to approximately only one fifth of the original sample size, but the two highest quintiles of risk both approximately reach significance at the 5% level in terms of absolute mortality reduction. When using reference class forecasting that interpolates between all risk quantiles, the trend between baseline risk and treatment effect becomes clearer. For example, panel B shows the variation in TATE when varying the distribution of baseline risk using exponential tilting. The estimation of ITEs using an Epanechnikov kernel with varying bandwidth gives very similar results (panel C2).

**Figure 2:**
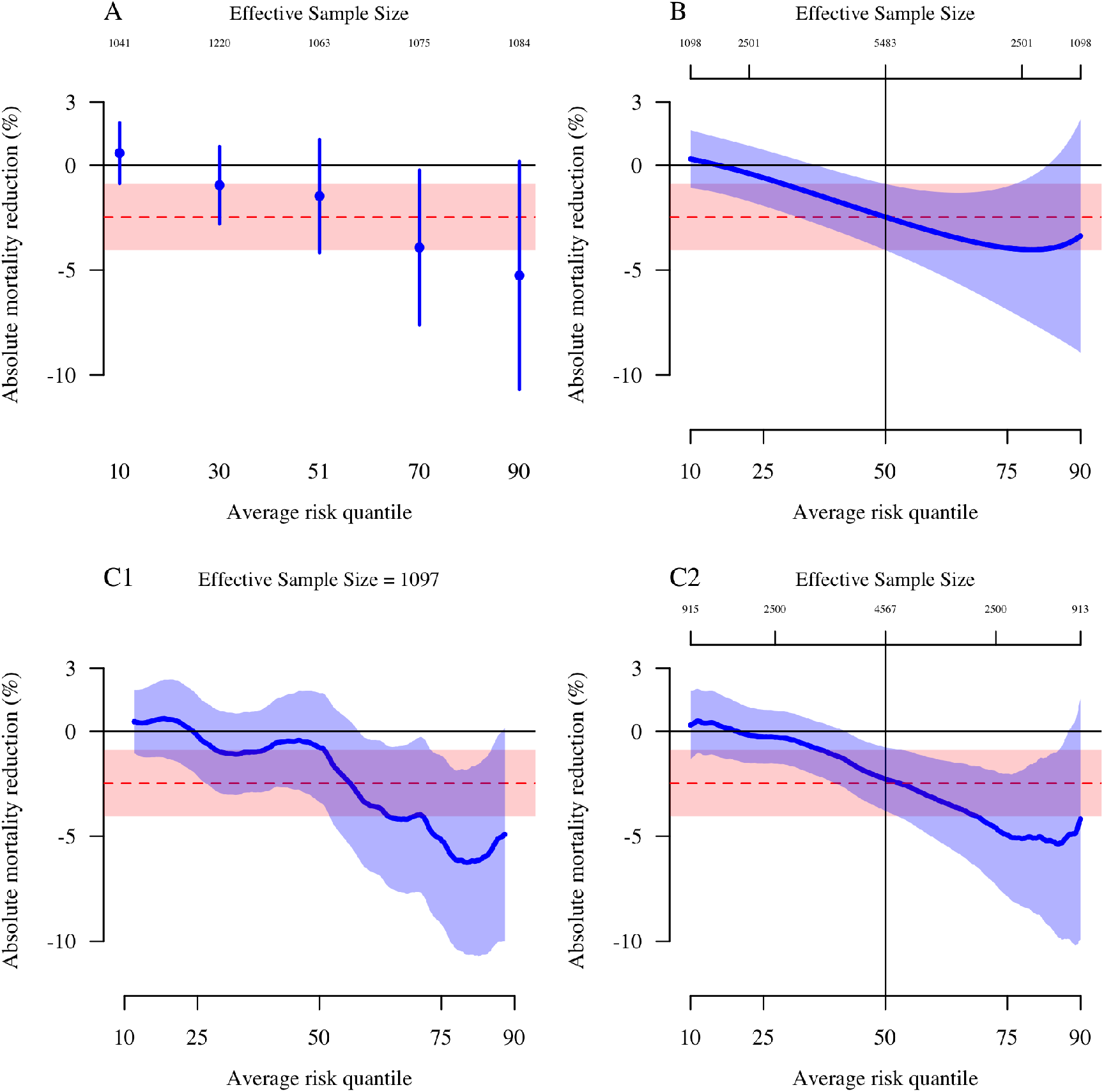
A graphical comparison of four approaches to reference class forecasting of ITEs (thick blue lines with 95% CIs shown as shaded blue areas) for patients enrolled in the AQUAMAT study [16]. In each panel the ATE (95% confidence interval) from the original trial (*n* = 5483) is shown by the dashed red line (red shaded area). The left column shows fixed bandwidth predictors (fixed effective sample sizes approximately equal to one fifth of the original sample size), and the right column shows varying bandwidth predictors (varying effective sample sizes). **A:** risk-based quintile partitioning. This does not interpolate between average risks in each subgroup. There is some minor variation in effective sample size due to ties in the multivariate risk scores. **B:** exponential tilting with free parameter *λ* as a global re-weighting scheme with varying effective sample size (top x-axis). This is centred around the overall treatment effect corresponding to the value *λ* = 0. **C1:** Epanechnikov kernel with fixed bandwidth chosen for an effective sample size of *n* = 1097 (20% of the original sample size). **C2:** Epanechnikov kernel with maximal bandwidth reference class. Note that the 50% risk quantile has an effective sample size reduction of 17% with respect to the original trial sample size due to the decay in weights. This contrasts with panel B where the ITE prediction at the 50% empirical risk quantile equals that of the ATE.

## Discussion

Randomised trials are designed and powered to estimate ATEs. However, the distribution of baseline risk is often highly skewed and the ATE will both overestimate and underestimate the benefit of treatment for patients who have lower or higher than average risk. A consequence of this variation in baseline risk and its influence on the reported treatment effect is that ATEs can be misleading when used to guide treatment recommendations at the individual level. It has been recommended that all trials report how treatment effect varies as a function of baseline risk, and the methods described here provide a principled framework for the graphical visualisation of HTEs and the estimation of ITEs and TATEs. This risk representation is an important example of a broader idea of constructing a reference class a priori on which to explore heterogeneity.

The classical approach to the estimation of ITEs is to construct a parametric model of the outcome conditional on each possible treatment assignment and observed key patient covariates. Typically this necessitates data transformations of the outcome measurement such as the log odds scale for logistic regression. Instead, we advocate a two-stage approach, first fitting a parametric model between the outcome and the key patient covariates preferably from an external data source: this is a multi-variate risk-model which does not involve counterfactuals. This risk model can then be used to construct causally valid weighted treatment effects which can be directly interpreted in terms in risk dependent ITEs and TATEs. In brief, instead of modelling the treatment effect directly, we recommend to model the risk and then use classical tests on randomised data to estimate ITEs. A major advantage of this approach is that it provides a formal approach to the use of prior observational data when evaluating the results of a randomised trial.

Graphical visualisation of RCT data using a well-defined risk-based reference class forecasting method such as exponential tilting of patients ‘information weight’ allows for a clear presentation of heterogeneity in treatment effect as a function of baseline risk. These reference class forecasting plots are defined on the scale of the original estimand and are explicitly centred around the ATE targeted by the original RCT, and explicitly show the loss in effective sample size as one attempts to predict in the tails of the risk distribution.

## Declarations

### Ethics approval and consent to participate

Not applicable to this study

### Consent for publication

Not applicable

### Availability of data and material

The data that support the findings of this study are available from the Mahidol Oxford Tropical Medicine Research Unit but restrictions apply to the availability of these data, which were used under license for the current study, and so are not publicly available. Data are however available from the authors upon reasonable request and with permission of the Mahidol Oxford Tropical Medicine Research Unit data-sharing committee.

### Authors’ contributions

JAW and CCH authors contributed equally to this work.

## Acknowledgements

We thank the Mahidol Oxford Tropical Medicine Research Unit for allowing us to use the data from the AQUAMAT study.

---

**Figure S1:**
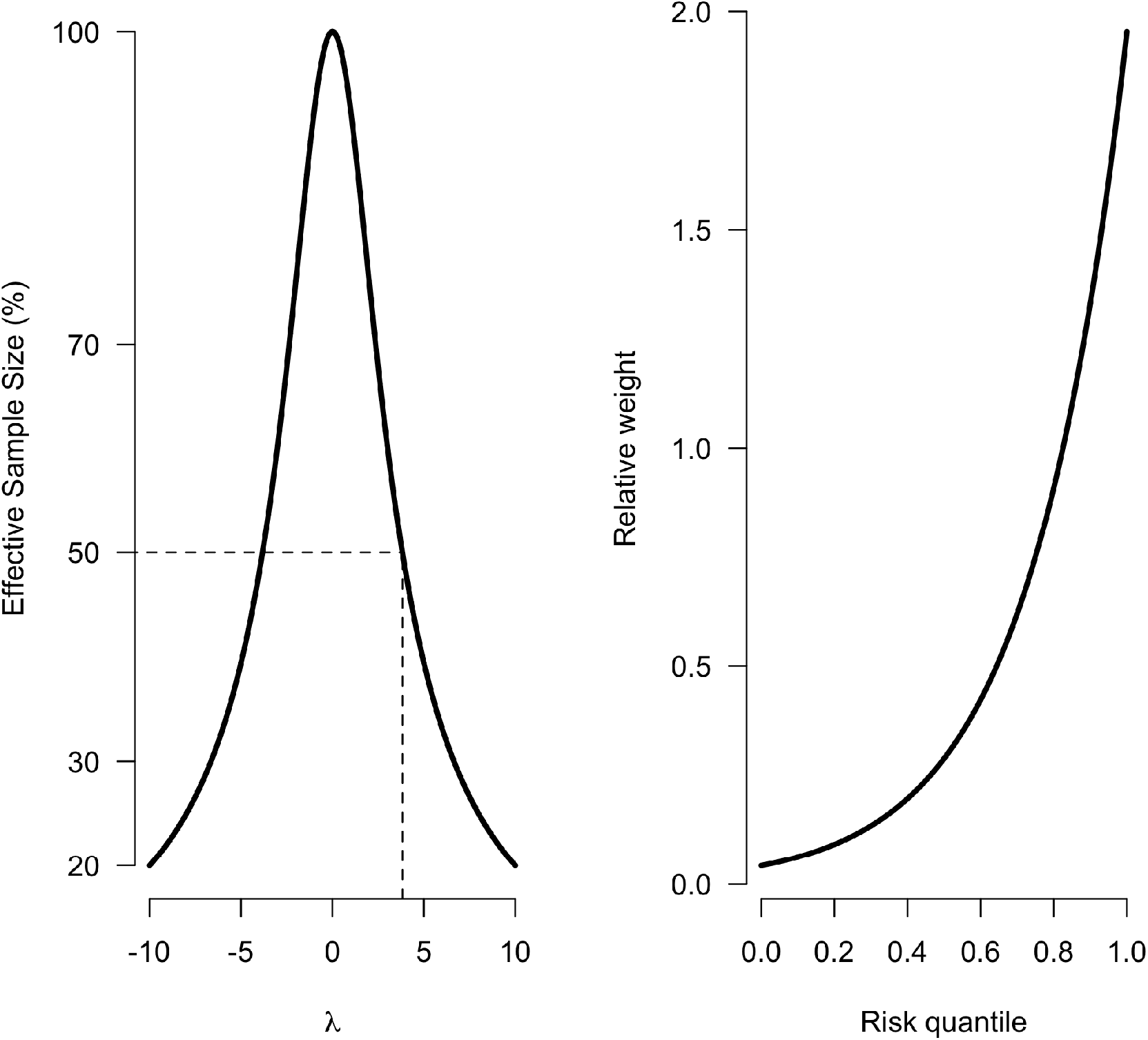
Exponential tilting of weights as a global reference class forecasting method. The left panel shows the relationship between the value of *λ* and the corresponding reduction in effective sample size (shown as a percentage of the original trial size). The right plot shows the distribution of weights according to risk for the value of *λ* (≈ 3.83) leading to a 50% reduction in effective sample size, tilting the risk distribution towards the higher risk patients.

**Figure S2:**
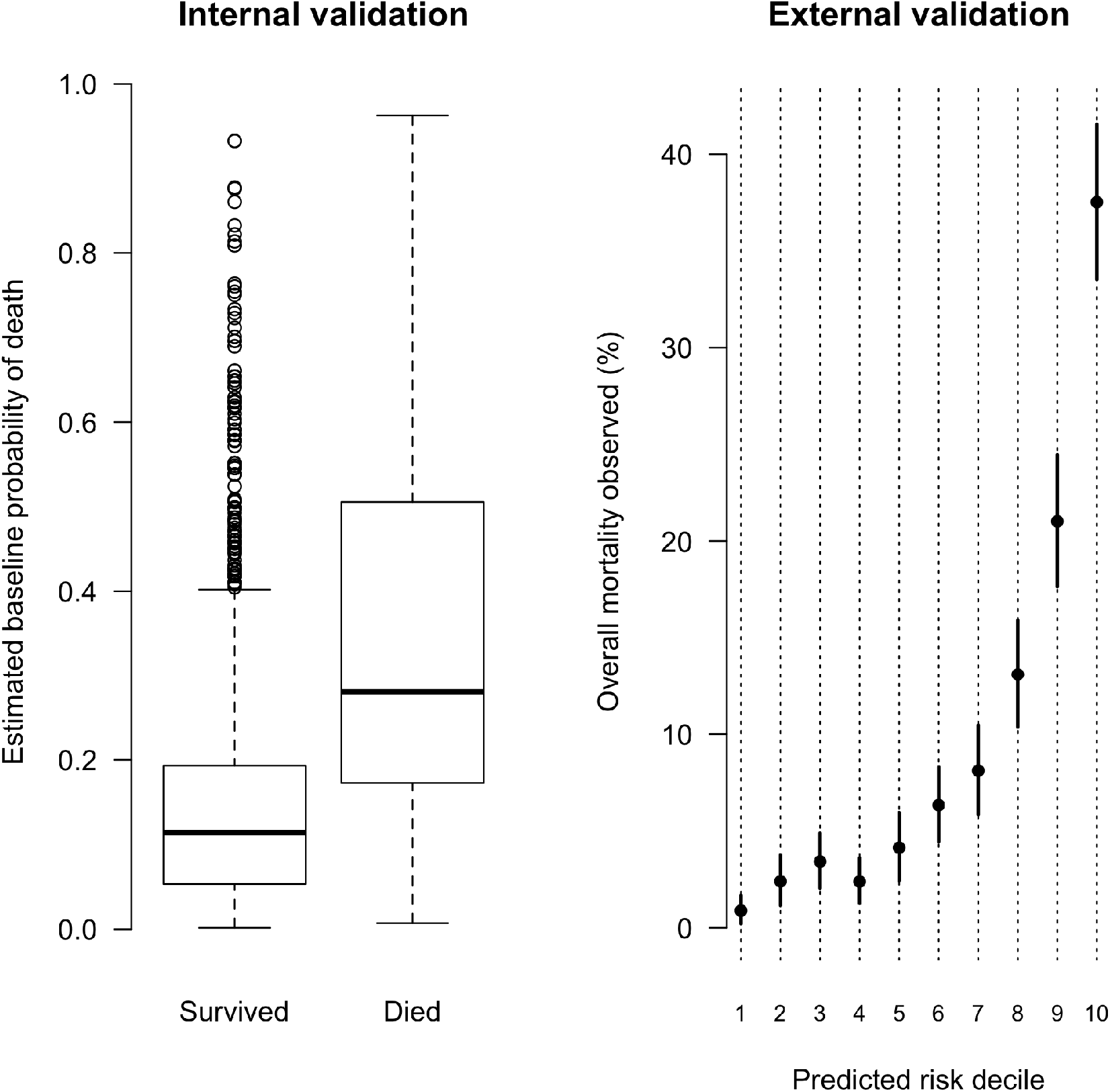
Internal (left) and external (right) of the multivariate risk model of death from severe malaria. The left plot shows a boxplot of estimated scores (probabilities of death from a logistic regression model) in patients who died versus those who survived. This is in the ‘training’ database consisting of a series of randomised and observational studies. The right plot shows the observed mortality in the AQUAMAT study as a function of predicted risk decile using this multivariate model.

**Figure S3:**
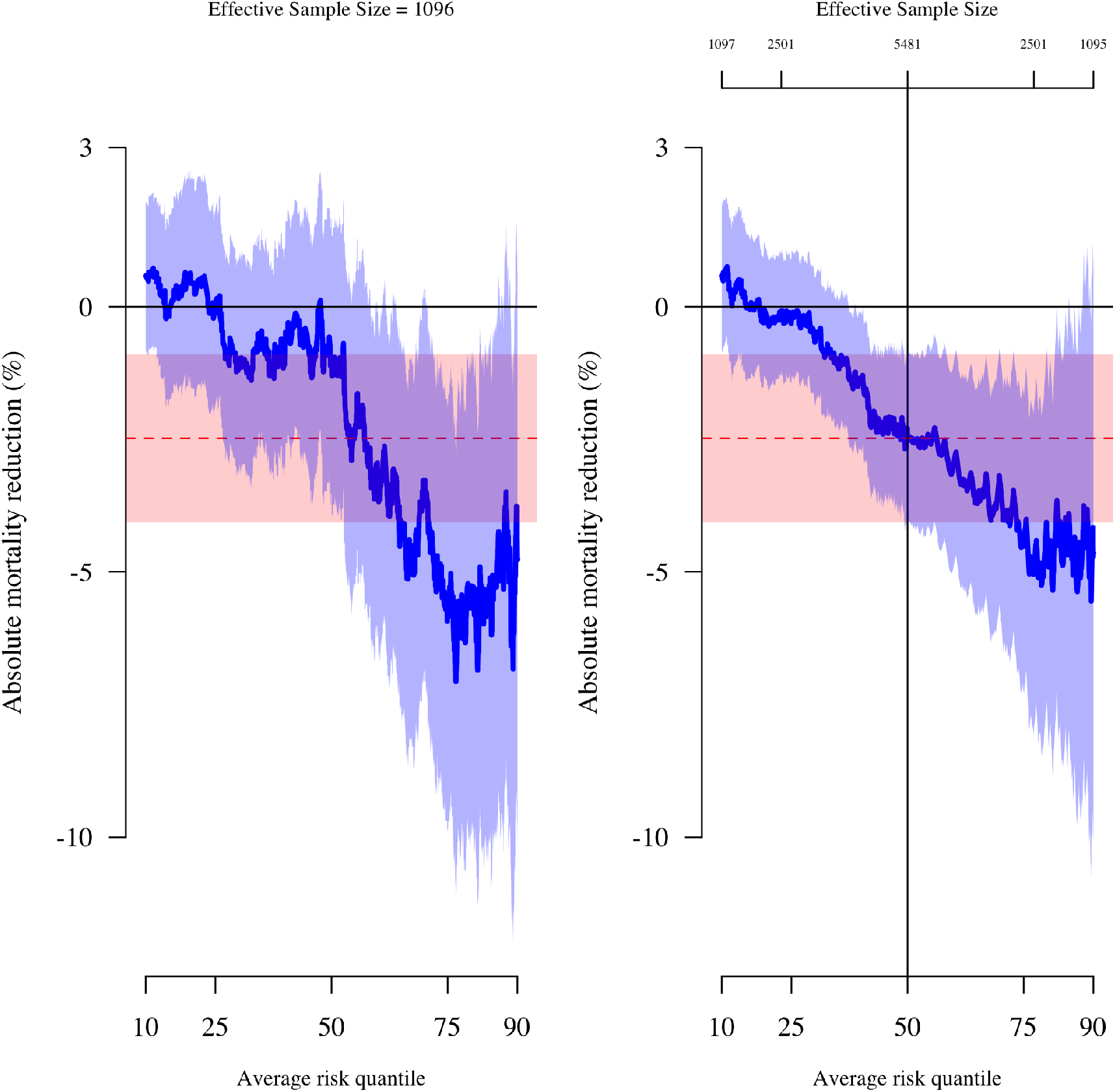
Boxcar kernel for reference class forecasting. This is fit to the same study data as in Figure 2. **Left**: boxcar kernel with fixed bandwidth. This corresponds to a *k*-nearest-neighbours approach to predicting the ITE with no smoothing. **right**: boxcar kernel with maximal bandwidth reference class.

## References

[1] David M Kent, Jason Nelson, Issa J Dahabreh, Peter M Rothwell, Douglas G Altman, and Rodney A Hayward. Risk and treatment effect heterogeneity: re-analysis of individual participant data from 32 large clinical trials. International journal of epidemiology, 45(6):2075–2088, 2016.

[2] David M Kent and Rodney A Hayward. Limitations of applying summary results of clinical trials to individual patients: the need for risk stratification. Jama, 298(10):1209–1212, 2007.

[3] David M Kent, Peter M Rothwell, John PA Ioannidis, Doug G Altman, and Rodney A Hayward. Assessing and reporting heterogeneity in treatment effects in clinical trials: a proposal. Trials, 11(1):85, 2010.

[4] David M Kent, Ewout Steyerberg, and David van Klaveren. Personalized evidence based medicine: predictive approaches to heterogeneous treatment effects. BMJ, 363:k4245, 2018.

[5] Jeremy B Sussman, David M Kent, Jason P Nelson, and Rodney A Hayward. Improving diabetes prevention with benefit based tailored treatment: risk based reanalysis of diabetes prevention program. Bmj, 350:h454, 2015.

[6] Xin Sun, John PA Ioannidis, Thomas Agoritsas, Ana C Alba, and Gordon Guyatt. How to use a subgroup analysis: usersâ [U+0080][U+0099] guide to the medical literature. Jama, 311(4):405–411, 2014.

[7] Peter M Rothwell. Subgroup analysis in randomised controlled trials: importance, indications, and interpretation. The Lancet, 365(9454):176–186, 2005.

[8] Matt P Wand and M Chris Jones. Kernel smoothing. Chapman and Hall/CRC, 1994.

[9] Elizbar A Nadaraya. On estimating regression. Theory of Probability & Its Applications, 9(1):141–142, 1964.

[10] Geoffrey S Watson. Smooth regression analysis. Sankhyā: The Indian Journal of Statistics, Series A, pages 359–372, 1964.

[11] James F Burke, Rodney A Hayward, Jason P Nelson, and David M Kent. Using internally developed risk models to assess heterogeneity in treatment effects in clinical trials. Circulation: Cardiovascular Quality and Outcomes, 7(1):163–169, 2014.

[12] Rodney A Hayward, David M Kent, Sandeep Vijan, and Timothy P Hofer. Multivariable risk prediction can greatly enhance the statistical power of clinical trial subgroup analysis. BMC medical research methodology, 6(1):18, 2006.

[13] Josh Hanson, Sue J Lee, Sanjib Mohanty, MA Faiz, Nicholas M Anstey, Prakay kaew Charun-watthana, Emran Bin Yunus, Saroj K Mishra, Emiliana Tjitra, Ric N Price, et al. A simple score to predict the outcome of severe malaria in adults. Clinical infectious diseases, 50(5): 679–685, 2010.

[14] WHO. Severe malaria. Tropical Medicine & International Health, 19(Supplement 1):7–131, 2014. doi: 10.1111/tmi.12313\_2. URL https://onlinelibrary.wiley.com/doi/abs/10.1111/tmi.12313_2.

[15] Arjen M Dondorp, François Nosten, Kasia Stepniewska, Nick Day, and Nick White. Artesunate versus quinine for treatment of severe falciparum malaria: a randomised trial. Lancet (London, England), 366(9487):717–725, 2005.

[16] Arjen M Dondorp, Caterina I Fanello, Ilse CE Hendriksen, Ermelinda Gomes, Amir Seni, Kajal D Chhaganlal, Kalifa Bojang, Rasaq Olaosebikan, Nkechinyere Anunobi, Kathryn Maitland, et al. Artesunate versus quinine in the treatment of severe falciparum malaria in african children (aquamat): an open-label, randomised trial. The Lancet, 376(9753):1647–1657, 2010.

[17] Elizabeth A Ashley, Mehul Dhorda, Rick M Fairhurst, Chanaki Amaratunga, Parath Lim, Seila Suon, Sokunthea Sreng, Jennifer M Anderson, Sivanna Mao, Baramey Sam, et al. Spread of artemisinin resistance in plasmodium falciparum malaria. New England Journal of Medicine, 371(5):411–423, 2014.

[18] Mallika Imwong, Tran T Hien, Nguyen T Thuy-Nhien, Arjen M Dondorp, and Nicholas J White. Spread of a single multidrug resistant malaria parasite lineage (pfpailin) to vietnam. The Lancet Infectious Diseases, 17(10):1022–1023, 2017.

[19] Richard J. Maude, Gofranul Hoque, Mahtab Uddin Hasan, Abu Sayeed, Shahena Akter, Rasheda Samad, Badrul Alam, Emran Bin Yunus, Ridwanur Rahman, Waliur Rahman, Romal Chowdhury, Tapan Seal, Prakaykaew Charunwatthana, Christina C. Chang, Nicholas J. White, M. Abul Faiz, Nicholas P.J. Day, Arjen M. Dondorp, and Amir Hossain. Timing of enteral feeding in cerebral malaria in Resource-Poor settings: A randomized trial. PLoS ONE, 6(11):1–7, 2011. ISSN 19326203. doi: 10.1371/journal.pone.0027273.

[20] Richard J Maude, Kamolrat Silamut, Katherine Plewes, Prakaykaew Charunwatthana, May Ho, M Abul Faiz, Ridwanur Rahman, Md Amir Hossain, Mahtab U Hassan, and Emran Bin Yunus. Randomized controlled trial of levamisole hydrochloride as adjunctive therapy in severe falciparum malaria with high parasitemia. The Journal of infectious diseases, 209(1): 120–129, 2013.

[21] Paul N Newton, Brian J Angus, Wirongrong Chierakul, Arjen Dondorp, Ronatrai Ruangveerayuth, Kamolrat Silamut, Pramote Teerapong, Yupin Suputtamongkol, Sornchai Looareesuwan, and Nicholas J White. Randomized comparison of artesunate and quinine in the treatment of severe falciparum malaria. Clinical infectious diseases, 37(1):7–16, 2003. ISSN 1537-6591. doi: 10.1086/375059.

[22] K Plewes, Hugh W. F. Kingston, Aniruddha Ghose, Thanaporn Wattanakul, Md. Mahtab Uddin Hassan, Md. Shafiul Haider, Prodip K. Dutta, Md. Akhterul Islam, Shamsul Alam, Selim Md. Jahangir, A. S. M. Zahed, Md. Abdus Sattar, M. A. Hassan Chowdhury, M. Trent Herdmen, Stije J. Leopold, Haruhiko Ishioka, Kim A. Piera, Prakaykaew Charunwatthana, Kamolrat Silamut, Tsin W. Yeo, Sue J. Lee, Mavuto Mukaka, Richard J Maude, Gareth D. H. Turner, Md. Abdul Faiz, Joel Tarning, John A. Oates, Nicholas M. Anstey, Nicholas J. White, Nicholas P. J. Day, Md. Amir Hossain, L. Jackson Roberts II, and Arjen M. Dondorp. Acetaminophen as a renoprotective adjunctive treatment in patients with severe and moderately severe falciparum malaria: a randomized, controlled, open-label trial. Clinical Infectious Diseases, (April), 2018. doi: 10.1093/cid/ciy213/4930781.

[23] Nguyen H Phu, Phung Q Tuan, Nicholas Day, Nguyen TH Mai, Tran TH Chau, Ly V Chuong, Dinh X Sinh, Nicholas J White, Jeremy Farrar, and Tran T Hien. Randomized controlled trial of artesunate or artemether in vietnamese adults with severe falciparum malaria. Malaria journal, 9(1):97, 2010.

[24] SEAQUAMAT group. Artesunate versus quinine for treatment of severe falciparum malaria: A randomised trial. Lancet, 366(9487):717–725, 2005. ISSN 01406736. doi: 10.1016/S0140-6736(05)67176-0.

[25] Tran Tinh Hien, Nicholas PJ Day, Nguyen Hoan Phu, Nguyen Thi Hoang Mai, Tran Thi Hong Chau, Pham Phu Loc, Dinh Xuan Sinh, Ly Van Chuong, Ha Vinh, Deborah Waller, et al. A controlled trial of artemether or quinine in vietnamese adults with severe falciparum malaria. New England Journal of Medicine, 335(2):76–83, 1996.

